## supplementary Fig 1-5 and Table1 for "Assembly of mTORC3 involves binding of ETV7 to two separate sequences in the mTOR kinase domain": ETV7-mTor_supplementary_materials 7-16-24.docx

**Supplemental Figures**

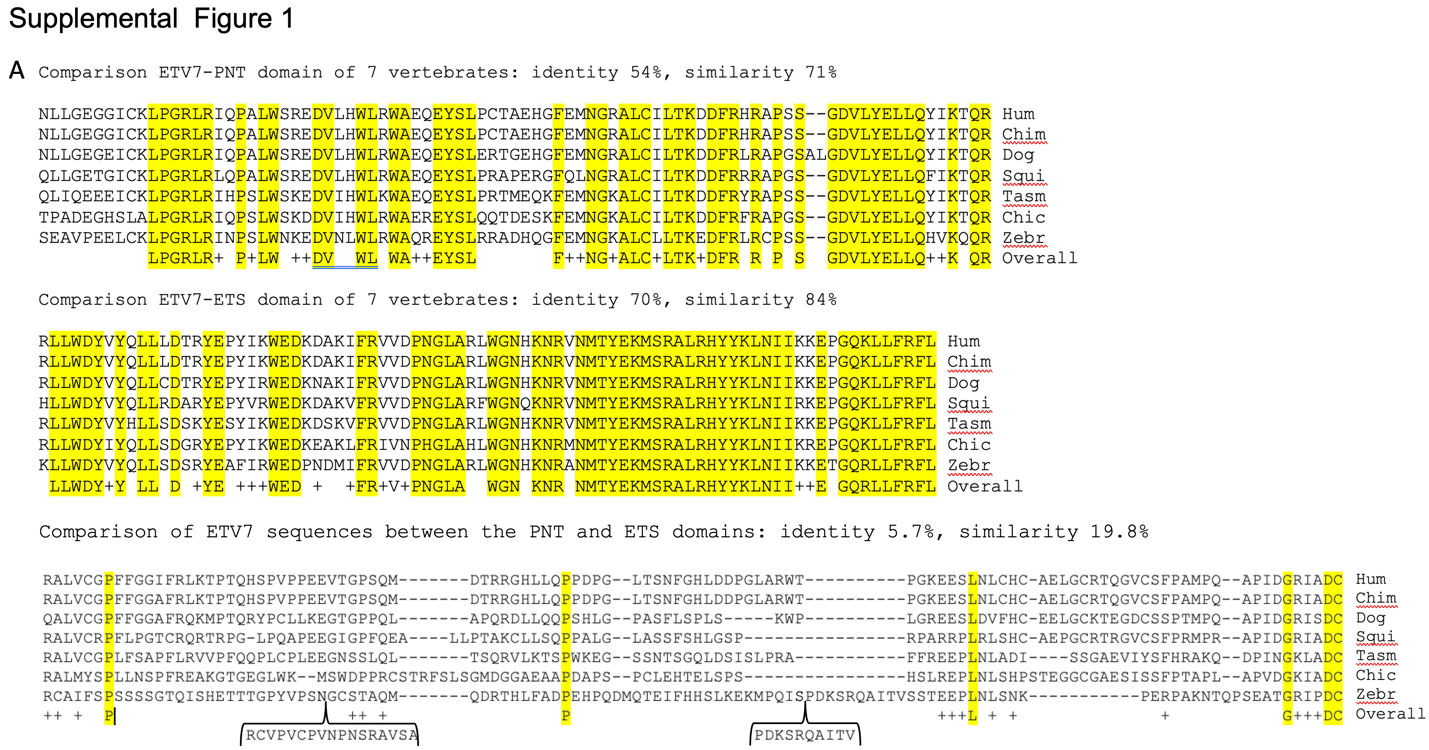

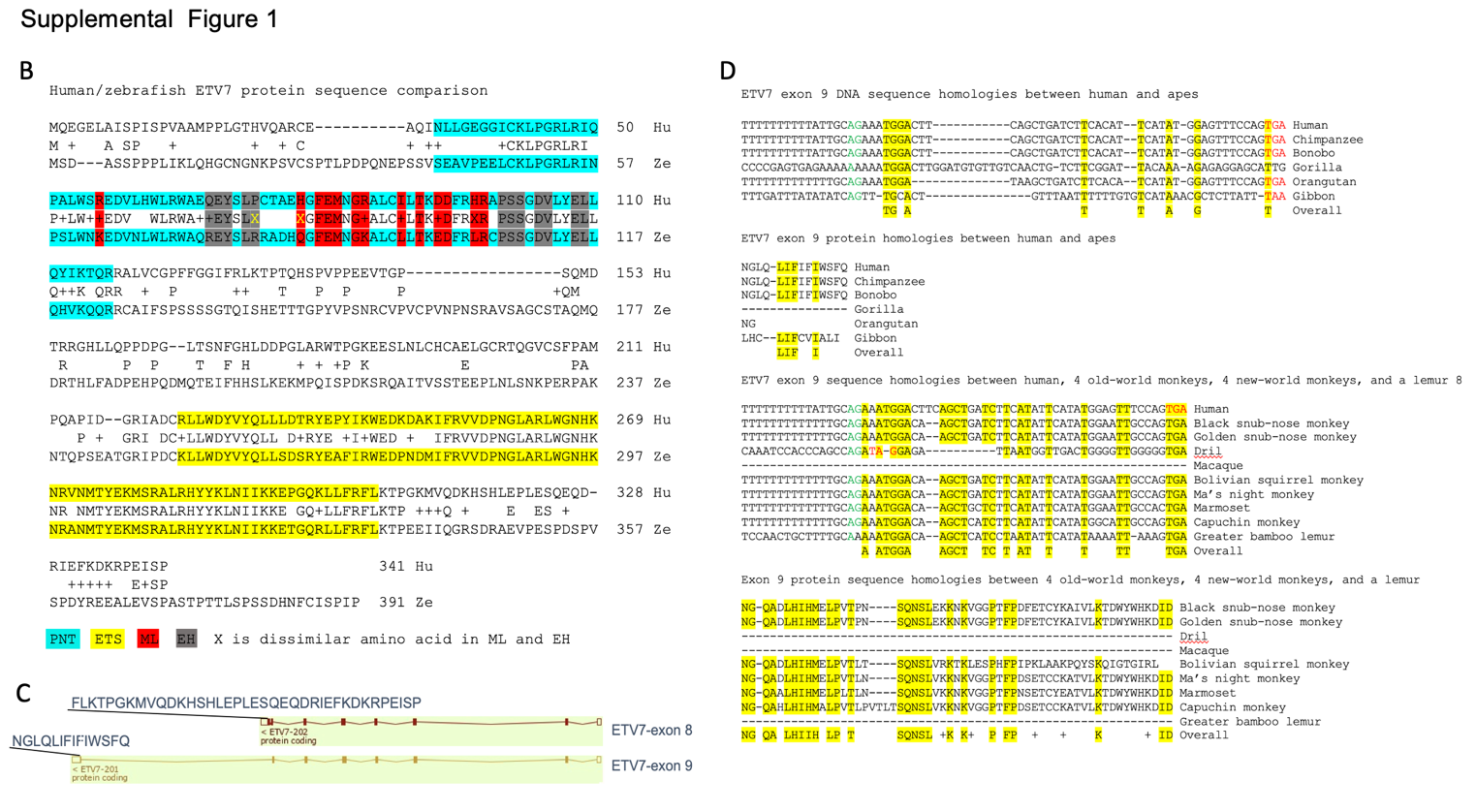

Supplementary Figure 1. Comparison of ETV7 sequences of different vertebrates.

1A. Comparison of ETV7 PNT, ETS, and sequences between the PNT and ETS domains of different vertebrates, including human (Hum), chimpanzee (Chim), dog (Dog), squirrel (Squi), Tasmanian devil (Tasm), chicken (Chick) and zebrafish (Zebr).

1B. Comparison of the ETV7 protein sequence of human (Hu) and zebrafish (Ze) ETV7. PNT sequences are highlighted in aquamarine, and ETS sequences are highlighted in yellow. Within the PNT domain, ML sequences are highlighted in red, and EH sequences in grey. X indicates dissimilar amino acids at identical positions in the ML and EH sequences of human and zebrafish.

1C. Genomic structure of ETV7-exon 8 and ETV7-exon 9 showing the protein sequence of the two dissimilar C-termini (https://useast.ensembl.org/).

1D. Comparison of exon 9 DNA and protein sequences of humans and apes and of humans, four old-world, four new-world monkeys, and a lemur. Identical base pairs or amino acids are highlighted in yellow, AG splice acceptor sequences in green, and stop codons in red. + indicates similar amino acids.

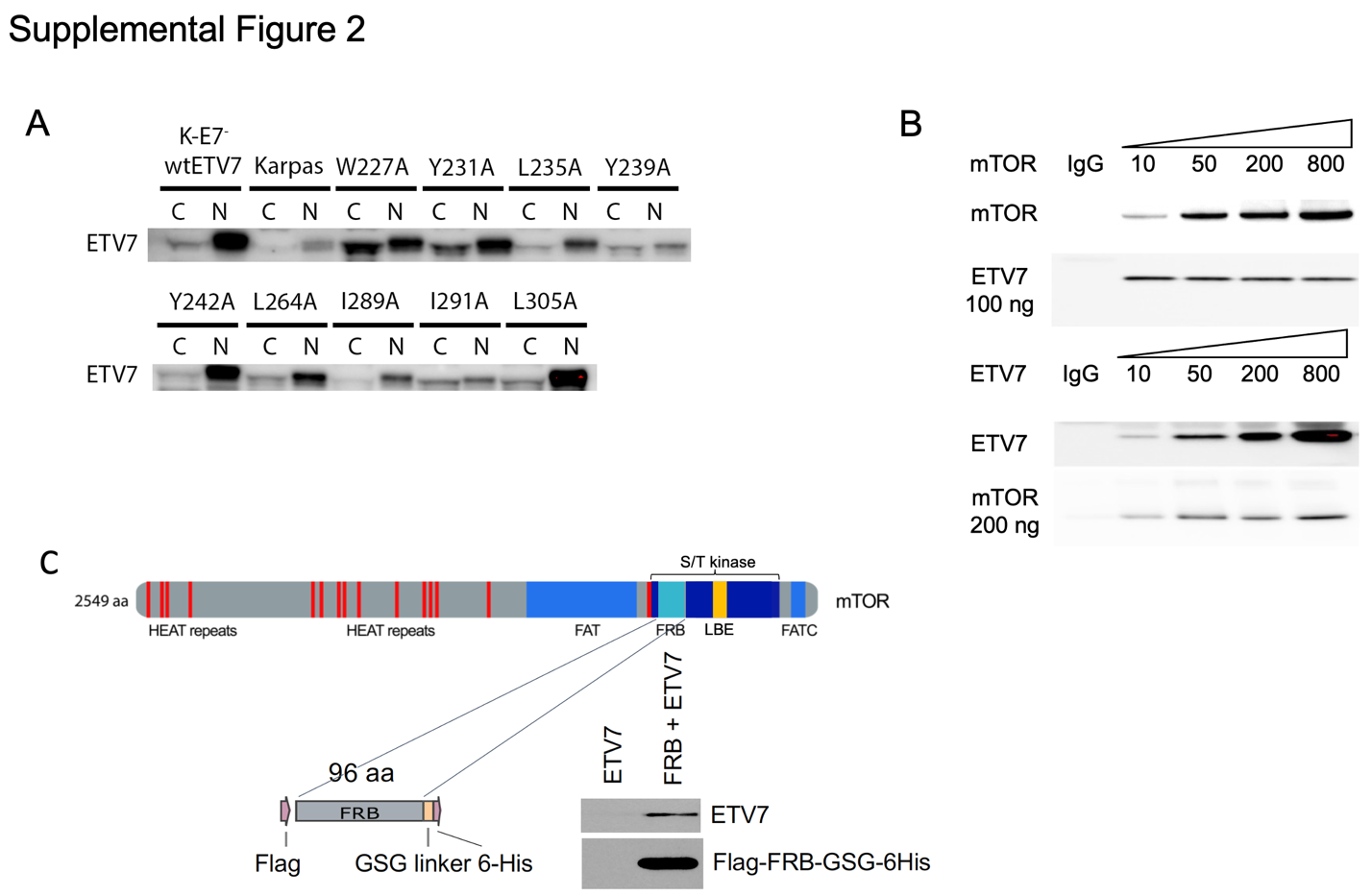

Supplementary Figure 2

Cellular localization of the ETV7 ETS mutants and ETV7-FRB binding *in vitro.*

2A. Western blot of cytoplasmic (C) and nuclear (N) fractions of KE7^-^ cells expressing wtETV7 (wild type), Karpas-299, or KE7^-^ cells expressing ETV7 carrying mutant ETS domains (W227A, Y231A, L235A, Y242A, L264A, I289A, I291A, L305A).

2B. *In vitro* association of purified mTOR and ETV7. Increasing amounts of mTOR were incubated overnight at 4 ° C with a steady amount of ETV7 (100 ng, top panel), and increasing amounts of ETV7 were incubated with a steady amount of mTOR (200 ng, bottom panel). Immunoblots of these ETV7 IPs (top panel) and mTOR IPs were probed for mTOR and ETV7.

2C. The top drawing shows the position of the 96 aa FRB fragment within the mTOR kinase domain (s/t kinase). Below is a blow-up drawing of the Flag-FRB-GSGlinker-6-His fragment, which was used in ETV7 binding experiments. On the right is an immunoblot of purified ETV7 and purified ETV7 + FRB protein, associated overnight at 4 ^0^C *in vitro,* and pulled down on Ni-NTA beads, probed for Flag and ETV7.

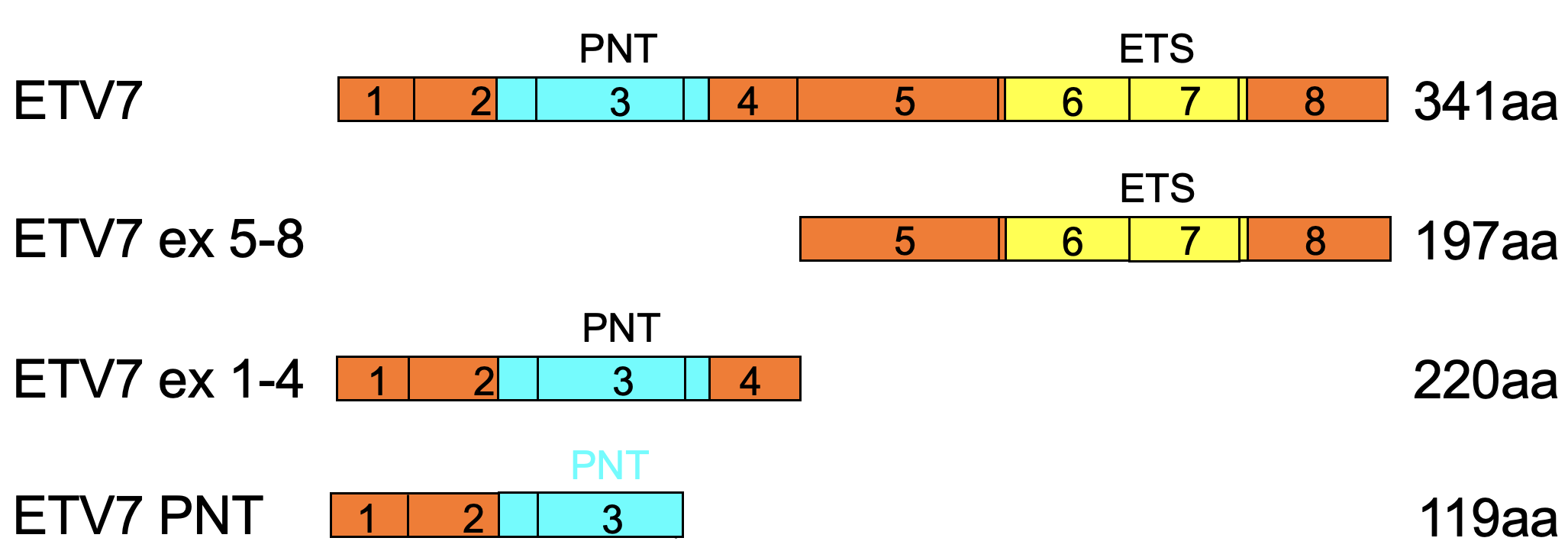

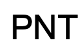

Supplemental Figure 3

Schematic showing different ETV7 deletions used for binding experiments with FRB and LBE protein fragments.

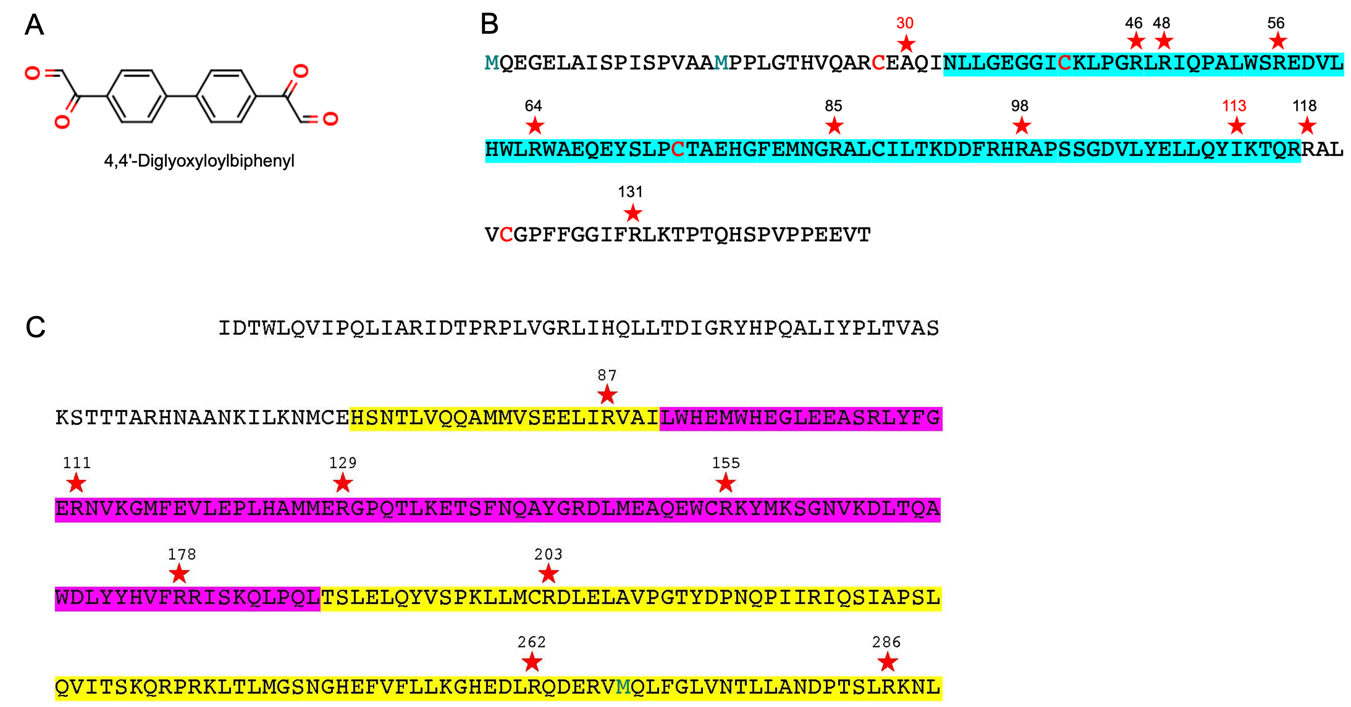

Supplemental Figure 4.

Chemical structure of the arginine cross-linker and cross-linked amino acids in the ETV7 PNT and mTOR FRB domains.

4A. Chemical structure of arginine crosslinker 4, 4'-diglyoxyloylbiphenyl.

4B. Position of cross-linked arginine amino acids in ETV7 marked by a star.

4C. Position of cross-linked amino acids in the FRB fragment of mTOR marked by a star.

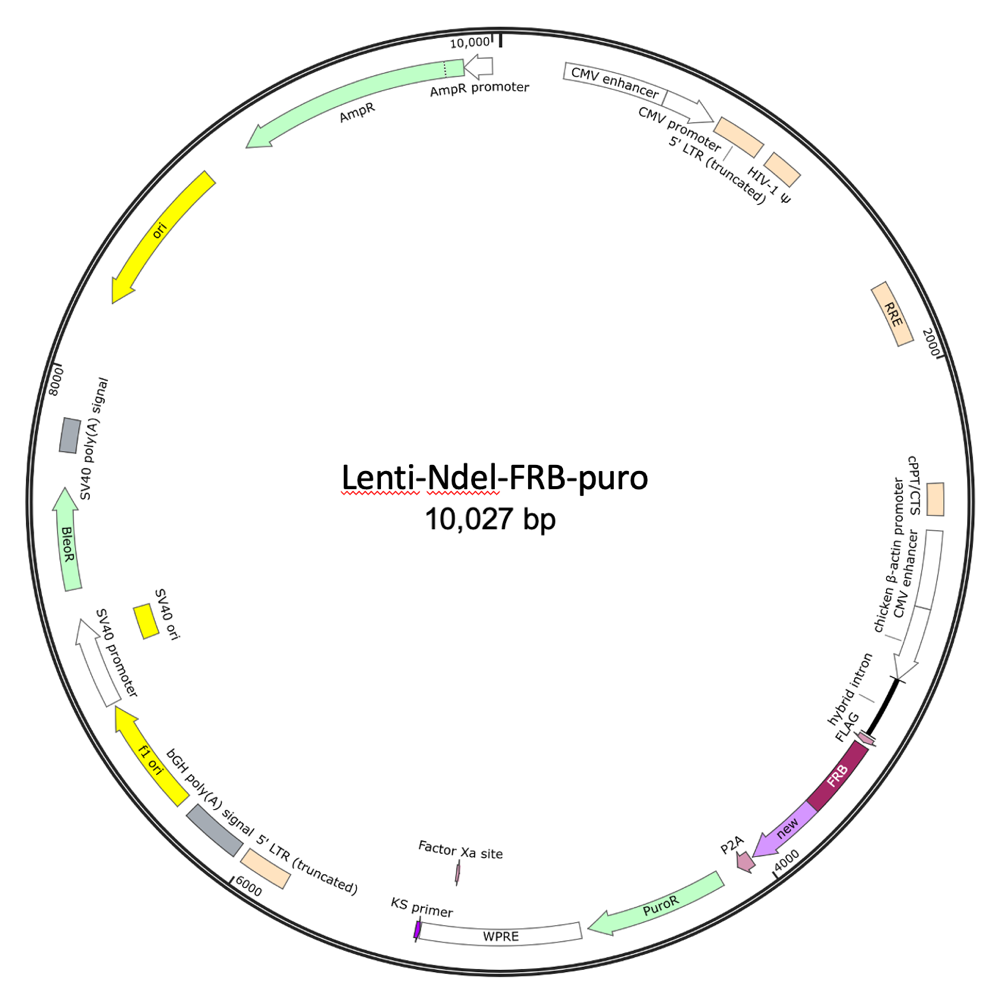

Supplemental Figure 5. Lenti-Ndel-FRB-puro.

Map of the lentiviral vector used for the expression of Ndel-FRB in Karpas-299 cells.

Supplemental Table 1. ΔΔG values for Alanine substitutions in the ETS domain of ETV7.

| **mutation** | **FoldX stability (kcal/mol)** | **ΔΔG (kcal/mol)** |
| --- | --- | --- |
| WT | -30.04 | - |
| W227 | -29.31 | 0.73 |
| Y231 | -28.88 | 1.16 |
| L235 | -29.29 | 0.75 |
| Y239* | -27.69 | 2.35 |
| Y242 | -27.63 | 2.41 |
| L264 | -28.12 | 1.92 |
| L289* | -29.29 | 0.75 |
| I291 | -28.26 | 1.78 |
| L305 | -29.06 | 0.98 |

*Alanine mutations that disrupt reciprocal co-IP

The stability of alanine mutants of the ETS domain of ETV7 as calculated by foldX with respect to the wild-type Alpha Fold model (Q9Y603), encompassing residues 223-305.
